# Molecular mapping of *YrTZ2*, a stripe rust resistance gene in wild emmer accession TZ-2 and its comparative analyses with *Aegilops tauschii*

**DOI:** 10.1101/131003

**Authors:** Zhenzhong Wang, Jingzhong Xie, Li Guo, Deyun Zhang, Genqiao Li, Tilin Fang, Yongxing Chen, Jun Li, Qiuhong Wu, Ping Lu, Yong Wang, Miaomiao Li, Haibin Wu, Yan Zhang, Wuyun Yang, Ming-Cheng Luo, Tzion Fahima, Zhiyong Liu

## Abstract

Wheat stripe rust, caused by *Puccinia striiformis* f. sp. *tritici* (*Pst*), is a devastating disease that can cause severe yield losses. Identification and utilization of stripe rust resistance genes are essential for effective breeding against the disease. Wild emmer accession TZ-2, originally collected from Mount Hermon, Israel, confers near-immunity resistance against several prevailing *Pst* races in China. A set of 200 F_6:7_ recombinant inbred lines (RILs) derived from a cross between susceptible durum wheat cultivar Langdon and TZ-2 was used for stripe rust evaluation. Genetic analysis indicated that the stripe rust resistance of TZ-2 to *Pst* race CYR34 was controlled by a single dominant gene, temporarily designated *YrTZ2*. Through bulked segregant analysis (BSA) and SSR mapping, *YrTZ2* was located on chromosome arm 1BS and flanked by SSR markers *Xwmc230* and *Xgwm413* with genetic distance of 0.8 cM (distal) and 0.3 cM (proximal), respectively. By applying wheat 90K iSelect SNP genotyping assay, 11 polymorphic loci (consist of 250 SNP markers) closely linked with *YrTZ2* were identified. *YrTZ2* was further delimited into a 0.8 cM genetic interval between SNP marker *IWB19368* and SSR marker *Xgwm413*, and co-segregated with SNP marker *IWB28744* (attached with 28 SNP markers). Comparative genomics analyses revealed high level of collinearity between the *YrTZ2* genomic region and the orthologous region of *Aegilops tauschii* 1DS. The genomic region between loci *IWB19368* and *IWB31649* harboring *YrTZ2* is orthologous to a 24.5 Mb genomic region between AT1D0112 and AT1D0150, spanning 15 contigs on chromosome 1DS. The genetic and comparative maps of *YrTZ2* provide framework for map-based cloning and marker-assisted selection (MAS) of *YrTZ2*.

## INTRODUCTION

Bread wheat (*Triticum aestivum* L.) is one of the top three most important food crops which production affects worldwide food security. Stripe rust, caused by *Puccinia striiformis* f. sp. *tritici* (*Pst*), is one of the severest wheat diseases worldwide. Growing resistance cultivars is the most cost-effective and environmentally friendly method to control this disease. Up to date, more than 60 formally named and many other provisionally designated stripe rust resistance genes or quantitative trait loci (QTL) have been reported (McIntosh *et al.* 2013, 2014, 2016). Out of them, two stripe rust resistance genes, *Yr36* and *Yr18*, have been isolated through map-based cloning strategy (Fu *et al.* 2009; Krattinger *et al.* 2009). However, stripe rust resistance genes tend to become ineffective due to the continuous evolution of *Pst* races and monoculture deployment of resistance cultivars in wide area (Wan *et al.* 2004, 2007; de Vallavieille-Pope *et al.* 2012). So there is a continued need for identification and utilization of diversified resistance genes from various wheat germplasm resources. Wild emmer (*Triticum turgidum* ssp. *dicoccoides*, 2n = 4x = 28, AABB) is allotetraploidy with A and B sub-genomes, which was derived from a spontaneous hybridization of two diploid wild grasses *Triticum urartu* (2n=14, AA) and an as yet unidentified *Aegilops* species related to *Aegilops speltoides* (2n=14, SS) (Dvorak *et al.* 1993; Dvorak and Zhang 1990). Wild emmer is the progenitor of cultivated tetraploid durum wheat (*Triticum turgidum* ssp. *durum*, AABB) and hexaploid bread wheat (*Triticum aestivum*, AABBDD) (Feldman 2001), harboring abundant genetic resources for wheat improvement, including abiotic stress tolerances (salt, drought and heat), biotic stress tolerances (powdery mildew, rusts and Fusarium head blight), grain protein quality and quantity, and micronutrient concentrations (Zn, Fe, and Mn) (Xie and Nevo 2008). Manual selection during wheat domestication resulted in an inadvertent loss of genes and quantitative trait loci (QTL) beneficial for improving wheat agronomic and economic traits. Although they could be introgressed into modern wheat cultivars through traditional long-term breeding methods, molecular breeding provides an improved strategy for wheat improvement and can greatly shorten the breeding period.

Previously, molecular markers used for genetic linkage maps are mainly comprised of restriction fragment length polymorphisms (RFLPs) (Blanco *et al.* 1998), amplified fragment length polymorphisms (AFLPs) (Nachit *et al.* 2001), simple sequence repeats (SSRs) (Somers *et al.* 2004; Song *et al.* 2005), and diversity arrays technology (DArT) (Akbari *et al.* 2006; Peleg *et al.* 2008). Recently, high-throughput genotyping technology became more and more important for genetic studies. With the advantages of abundance, usual biallelism and availability of genotyping platform, single nucleotide polymorphisms (SNPs) are increasingly applied for high-density genetic mapping, physical map construction, comparative genomics analysis, genome-wide association studies (GWAS) and genomic selection in rice (Zhao *et al.* 2011), maize (Ganal *et al.* 2011; Riedelsheimer *et al.* 2012), and wheat (Akhunov *et al.* 2009; Luo *et al.* 2009; Cavanagh *et al.* 2013; Wang *et al.* 2014).

Fine mapping and map-based cloning of wheat genes is tedious because of the characteristics of wheat genome: allopolyploid (AABBDD), large genome size (17 gigabase) and numerous repeat DNA (90%). The availability of draft genome sequences and International Wheat Genome Sequencing Consortium (IWGSC) survey sequences of *T. aestivum* cv. Chinese Spring, *T. urartu* accession G1812 and *Ae. tauschii* accession AL8/78 (Brenchley *et al.* 2012; Jia *et al.* 2013; Ling *et al.* 2013; IWGSC 2014) facilitates wheat gene mapping. In particular, the released high-resolution SNP genetic linkage map and accurate physical map of *Ae. tauschii* accession AL8/78 provides closely wheat-related target for comparative genomics analyses (Luo *et al.* 2013).

In present study, we report: (1) the identification and genetic mapping a near-immunity stripe rust resistance gene *YrTZ2* derived from wild emmer with microsatellite markers and 90K iSelect SNP genotyping assay, and (2) comparative genomics analysis of the genomic regions of *YrTZ2* with the genetic linkage map and physical map of *Ae. tauschii*.

## MATERIALS AND METHODS

### Plant Material

The wild emmer accession TZ-2 was used as the stripe rust resistant parent to make cross with a highly susceptible durum wheat cultivar Langdon. A set of 200 F_6:7_ recombinant inbred lines (RILs) advanced by single-seed descent approach and the parental Langdon and TZ-2 were evaluated for stripe rust resistance with the prevailing *Pst* race CYR34. A highly susceptible wheat variety Mingxian169 was used as the susceptible control.

### Stripe rust evaluations

The parental lines Langdon and TZ-2, Langdon/TZ-2 hybrid F_1_, 200 F_6:7_ RILs and susceptible control Mingxian169 were inoculated with *Pst* race CYR34 at the jointing stage in Chengdu of Sichuan Province, China. At 18 - 20 days post inoculation when the susceptible control Mingxian169 had become severely infected, the infection type (IT) was recorded with a scale of 0 - 4, with 0 (immune reaction), 0; (hypersensitive reaction), 1 (highly resistant), 2 (moderately resistant), 3 (moderately susceptible) and 4 (highly susceptible), the values of 0 - 2 were rated as resistant, and those of 3 - 4 were rated as susceptible (Zhang *et al.* 2001). ITs were recorded again ten days later.

### Genomic DNA isolation and SSR marker analysis

Genomic DNAs of the parental lines and the F_6:7_ RILs population were extracted from seeding leaves using the Plant Genomic DNA Kit (Tiangen Biotech, CO., Ltd, Beijing, China). DNA concentration was quantified using NanoPhotometer® P360 (Implem GmbH, Munich, Germany) and normalized to 100 ng/ul. Resistant and susceptible DNA bulks were produced by separately mixing equal amounts of DNA from ten homozygous resistant and ten homozygous susceptible F_6:7_ families for bulked segregant analysis (Michelmore *et al.* 1991). Wheat genomic SSRs (*Xgwm*, *Xwmc*, *Xbarc*, *Xcfa*, and *Xcfd* series, https://wheat.pw.usda.gov) were used for polymorphism surveys between the two DNA bulks, and the polymorphic SSR markers were subsequently genotyped in the RIL mapping populations.

PCR reactions were carried out in a 10 μl reaction volume with the following conditions: one denaturation cycle at 94° for 5 min, followed by 35 cycles at 94° for 45 s, 55 - 65° (depending on specific primers) for 45 s, and 72° for 1 min, followed by an extension step of 72° for 10 min. Fragment analysis of PCR products were carried out on 8% non-denaturing polyacrylamide gels (39 acrylamide: 1 bisacrylamide). After electrophoresis, the gels were silver stained and photographed.

### Infinium 90K iSelect SNP Genotyping

To saturate the genomic region harboring the stripe rust resistance gene, the 200 F_6:7_ RILs were genotyped using wheat 90K iSelect SNP genotyping assay platform at the Genome Center of University of California, Davis according to the manufacturer’s protocol. SNP allele clustering was conducted with two population-based detection algorithms: Density Based Spatial Clustering of Applications with Noise (DBSCAN) and Ordering Points to Identify the Clustering Structure (OPTICS) using the polyploid version of GenomeStudio software as described in Wang *et al.* (2014). Subsequently, the cluster matrix of polymorphic SNP markers was output from the polyploid version of GenomeStudio, and the genotypes of samples assigned in TZ-2 cluster were marked ‘1’, and the genotypes of sample located in Langdon cluster were marked ‘2’, the others were marked ‘0’.

### Genetic mapping of the stripe rust resistance gene

The polymorphic SNP markers, SSR markers and stripe rust resistance genotypes were used for linkage analysis with the MultiPoint mapping software as described in Peleg *et al.* (2008) and Luo *et al.* (2013). Co-segregating SNP markers were regarded as a polymorphic locus. The linkage map was constructed with the software Mapdraw V2.1 (Liu and Meng 2003).

### Data availability

The authors state that all data necessary for confirming the conclusions presented in the article are represented fully within the article.

## RESULTS

### Inheritance of the stripe rust resistance gene in TZ-2

The wild emmer accession TZ-2 and durum wheat cultivar Langdon showed nearly immune and highly susceptible to stripe rust, respectively. The F_1_ plants are highly resistant to CYR34, indicating the dominant nature of the stripe rust resistance in TZ-2. Of the 200 F_6:7_ RILs derived from the cross between Langdon and TZ-2, 103 were resistant (IT 0-2) and 97 were susceptible (3-4), which fits the expected 1:1 ratio for a single gene inheritance (χ^2^_1:1_= 0.18, P<0.05), indicating that a single dominant locus, provisionally designated *YrTZ2*, in TZ-2 is responsible for the stripe rust resistance.

### Identification of microsatellite markers linked to *YrTZ2*

Initially, 194 SSR primer pairs distributed randomly throughout the whole genome were screened for polymorphisms between the parental lines as well as the resistant and susceptible DNA bulks. SSR markers, *Xwmc406*, *Xwmc230*, *Xgwm413*, *Xwmc128* and *Xcfd65* revealed polymorphisms between the resistant and susceptible parents as well as the bulked segregants. After testing the F_6:7_ segregating population, a linkage map for stripe rust disease resistance gene *YrTZ2* was constructed. The gene *YrTZ2* was localized into a 1.1 cM genetic interval between SSR markers *Xwmc230* and *Xgwm413* (Fig. 1).

**Figure 1.**
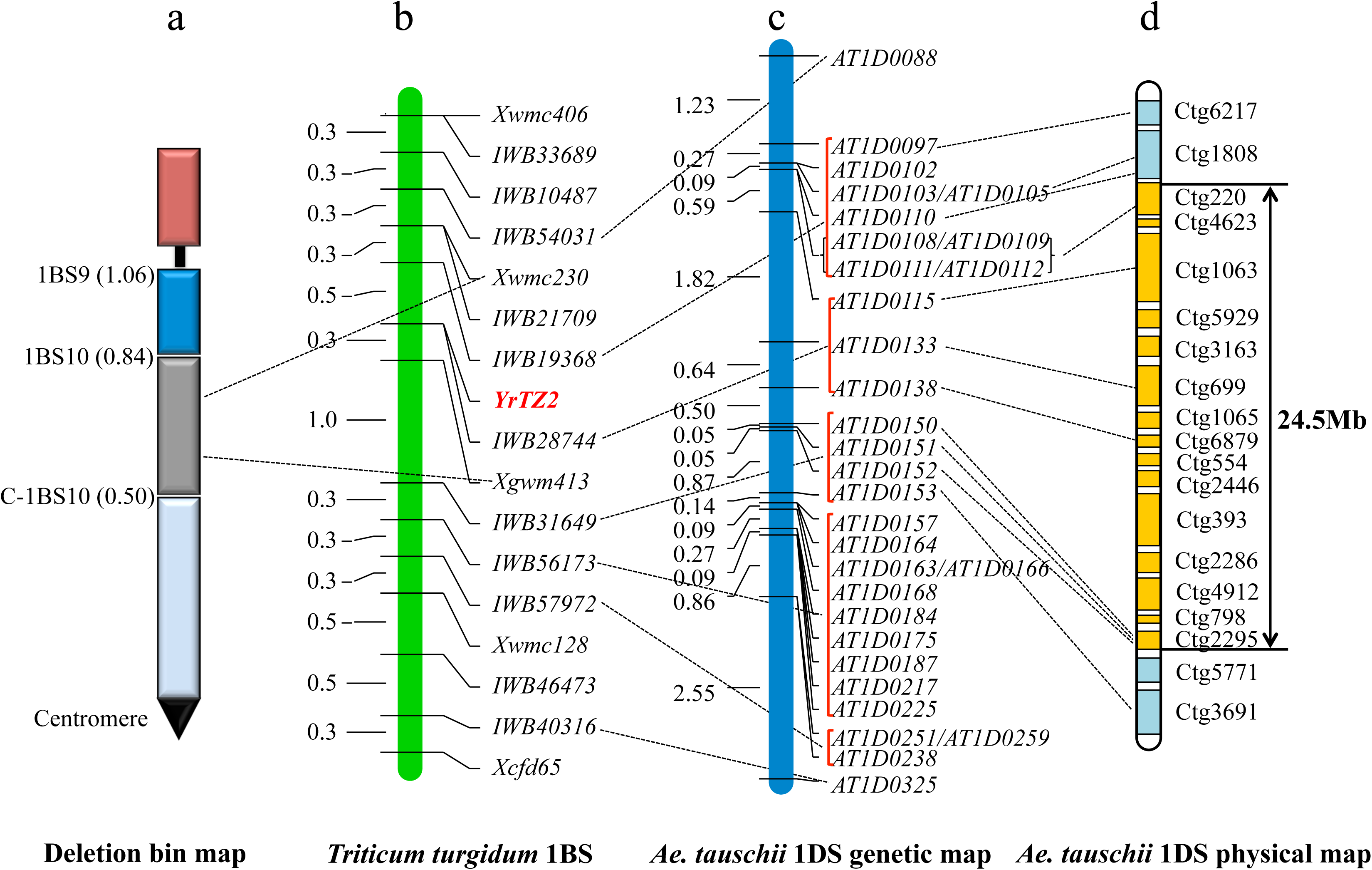
Genetic linkage map of the stripe rust resistance gene *YrTZ2*

### Chromosome arm assignment and physical bin mapping

In order to locate the *YrTZ2* in the deletion bins on chromosome 1BS, Chinese Spring homoeologous group 1 nullisomic-tetrasomics, ditelosomics and deletion lines were used to assign the chromosomal and physical bin locations of the *YrTZ2-*linked SSR markers. Both SSR markers *Xgwm413* and *Xwmc230* were detected in N1A-T1B, N1D-T1A, Dt1BS and 1BS-9, but absent in N1B-T1A, Dt1BL and 1BS-10 (Fig. 2), indicated that *YrTZ2* is located on chromosome 1BS bin 0.50-0.84 (Fig. 1).

**Figure 2.**
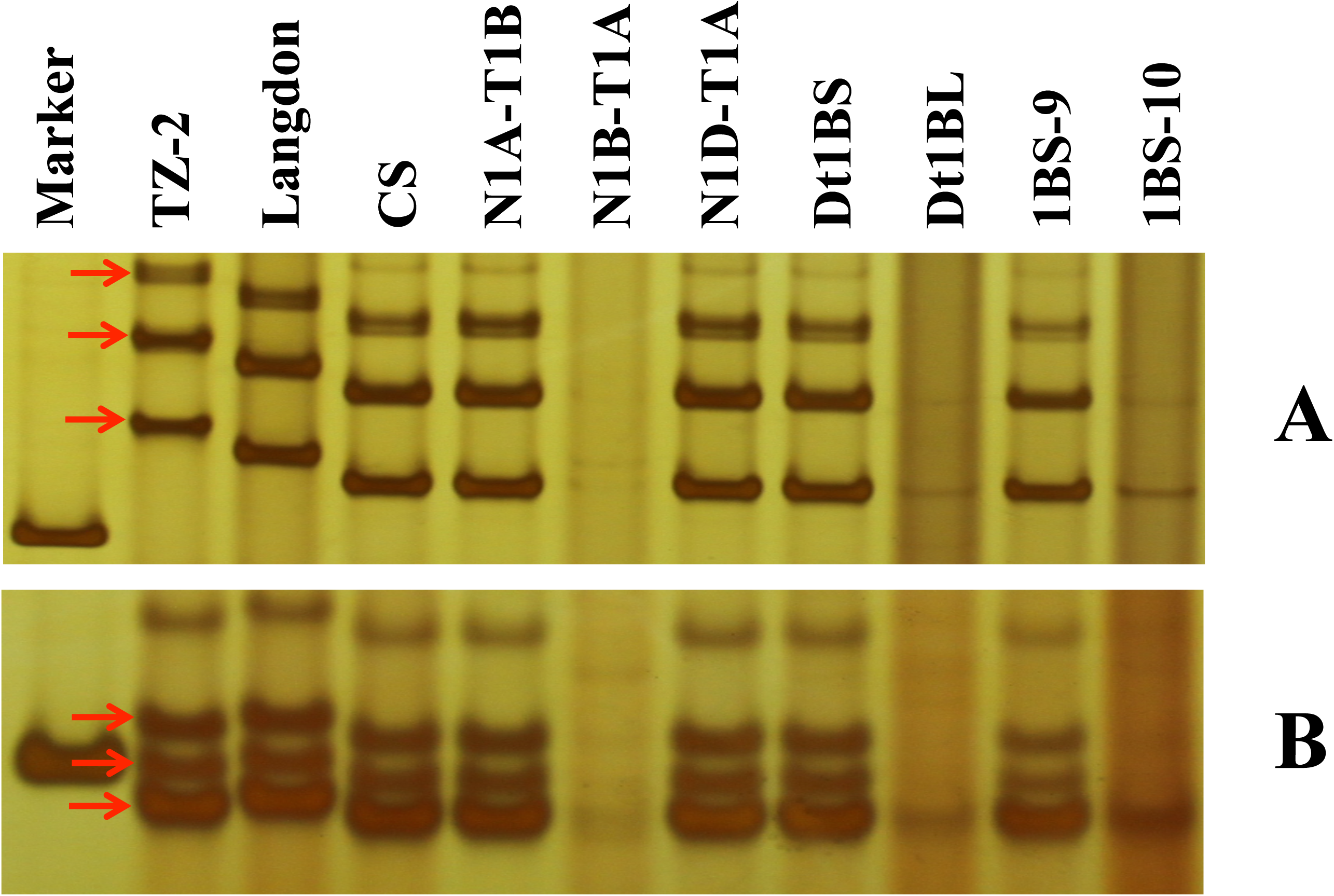
Amplification patterns of markers *Xwmc230* (**2A**) and *Xgwm413* (**2B**) in the parental lines TZ-2 and Langdon, Chinese Spring (CS) and its homoeologous group 1 nullisomic - tetrasomics, ditelosomics, and deletion lines

### Identification of SNP markers linked to *YrTZ2*

The 200 F_6:7_ RILs were genotyped with 90K iSelect SNP genotyping assay. After clustering, 15625 SNP markers were polymorphic between the parental lines, which were subsequently used for whole genome linkage map construction utilizing the MultiPoint mapping software. Polymorphic SNP and SSR markers linked to *YrTZ2* were used to construct a high-resolution linkage map of *YrTZ2.* Due to the limitation of population size, multiple co-segregating SNP markers were attached to one polymorphic locus with minimum missing scores and used as skeleton marker. All together, 11 polymorphic loci (consisting of 250 SNP markers), *IWB33689*, *IWB10487*, *IWB54031*, *IWB21709*, *IWB19368*, *IWB28744*, *IWB31649*, *IWB56173*, *IWB57972*, *IWB46473* and *IWB40316*, were integrated into the genetic linkage map of *YrTZ2* (Fig. 1). *YrTZ2* was finally delimited into an 0.8 cM interval between SNP locus *IWB19368* and SSR marker *Xgwm413*, and co-segregated with SNP locus *IWB28744* (attached with 28 SNP markers) (Fig. 1).

### Identification collinearity genomic region of *YrTZ2* in *Ae*. *tauschii* and comparative genomics analysis

The sequences of the 250 SNP markers clustered into 11 polymorphic loci were used as queries to search the *Ae. tauschii* SNP marker extended sequence database to identify the orthologous gene pairs between *T. dicoccoides* 1BS and *Ae. tauschii* 1DS. Out of the 11 polymorphic loci, 7 loci, *IWB54031*, *IWB19368, IWB28744, IWB31649, IWB56173, IWB57972* and *IWB40316*, identified 31 orthologous SNP marker extended sequence in *Ae. tauschii*. Compararive genomics analysis revealed high levels collinearity between *YrTZ2* genomic region and its orthologous genomic regions in *Ae. tauschii* 1DS (Fig. 1; Table S1).

*YrTZ2* was mapped between SNP markers *IWB19368* and *IWB31649*, and co-segregated with *IWB28744*. *IWB19368* and *IWB31649* are corresponding to the extended sequences of markers AT1D0112 (distal) and AT1D0150 (proximal), respectively, on chromosome 1DS that were anchored to the assembled BAC contigs ctg220 and ctg2295 in the physical map of *Ae*. *tauschii*. Therefore, the genomic region between *IWB19368* and *IWB31649* was orthologous to a 24.5Mb genomic region containing 15 BAC contigs, ctg220, ctg4623, ctg1063, ctg5929, ctg3163, ctg699, ctg1065, ctg6879, ctg554, ctg2446, ctg393, ctg2286, ctg4912, ctg798 and ctg2295 on chromosome 1DS (Fig. 1).

## DISCUSSION

Bread wheat is serving as an important global food crop all the time. Maximizing wheat production is becoming a big challenge for researchers, breeders and growers. Wild relatives of wheat harbor rich genetic resource for wheat improvement (Schneider *et al.* 2008; Xie and Nevo 2008). Wild emmer is the ancestor of modern cultivated wheat and mainly distributed in central-eastern (Turkey, Iran and Iraq) and western areas (Syria, Lebanon, Jordan and Israel) of the Fertile Crescent (Avni *et al.* 2014). Wild emmer harbors abundant beneficial traits that can be introgressed into tetraploid and hexaploid wheat in modern wheat breeding programs. However, wild emmer has not been explored thoroughly and its potential in wheat breeding programs remains to be further characterized (Xie and Nevo 2008).

Wild emmer accession TZ-2 was collected from Mount Hermon, Israel, and showed highly stripe rust resistance to many *Pst* races (CYR29, CYR30, CYR31, CYR32, CYR33 and CYR34) in greenhouse seedling and field adult plant stage tests. In this study, genetic analysis showed that the stripe rust resistance to CYR34 in TZ-2 is controlled by a single dominant gene *YrTZ2* that was mapped between SNP locus *IWB19368* and SSR marker *Xgwm413* in a 0.8 cM genetic interval on chromosome 1BS Bin 0.50-0.84. Up to date, another two stripe rust resistance genes, *Yr15* and *YrH52*, were derived from Israeli wild emmer wheat and located on chromosome 1BS. *Yr15* was identified from wild emmer accession G25 and mapped on chromosome 1BS using cytogenetic analysis (McIntosh *et al.* 1996) and molecular markers (Sun *et al.* 1997; Chagué *et al.* 1999; Ramirez-Gonzalez *et al.* 2015). When studying the stripe rust resistance gene in *T. dicoccoides* accession Hermon 52, Peng *et al.* (1999, 2000) found the resistance locus *YrH52* was linked to SSR marker *Xgwm413* with a genetic distance of 1.3 cM (proximal). *YrH52*-linked polymorphic microsatellite markers analysis revealed that *Yr15* (*Xgwm413/UBC212a*-*Yr15*-*Nor1*) is different from *YrH52* (*Xgwm413/UBC212a/Nor1*-*YrH52*-*Xgwm273*) on 1BS (Peng *et al.* 2000). In current study, *YrTZ2* (*Xgwm413-YrTZ2-IWB19368*) was located at similar portion of chromosome 1BS as that of *Yr15* and *YrH52*. Phytopathology and allelism tests need to be conducted in the future to clarify if *YrTZ2* is allelic or closely linked to *Yr15* or *YrH52*.

In addition to *Yr15*, *YrH52* and *YrTZ2*, several other stripe rust resistance genes have been identified on chromosome 1BS. *Yr10* was identified from Turkish hexaploid wheat accession PI 178383 and mapped at the terminal region of chromosome 1BS (Wang *et al.* 2002). *Yr24* was derived from *T*. *turgidum* subsp. *durum* accession K733 (McIntosh and Lagudah 2000). *Yr26* was assumed to be from durum line γ80-1, a γ-radiated mutant (Ma *et al.* 2001). *YrCH42* was identified from Chinese wheat cultivar Chuanmai 42 (Li *et al.* 2006). Evidences showed that *Yr24*, *Yr26* and *YrCH42* were the same gene (Ma *et al.* 2001; Li *et al.* 2006; McIntosh *et al.* 2013) and is losing resistance to the new virulent *Pst* race CYR34 in China (Han *et al.* 2012). *YrAlp* was derived from spring wheat cultivar Alpowa with race-specific all-stage resistance (Lin and Chen 2007). Cheng *et al.* (2014) identified broad-spectrum all-stage stripe rust resistance genes *Yr64* and *Yr65* in different bins of chromosome 1BS from durum wheat accessions PI 331260 and PI 480016, respectively. Pyramiding these genes on chromosome 1BS via marker-assisted selection would benefit the development of durable and broad-spectrum stripe rust resistance varieties in wheat breeding program. The characteristics of large genome size, hexaploid nature and numerous repetitive DNA sequences presented a formidable challenge to fine mapping and map-based cloning of wheat genes. Single nucleotide polymorphisms (SNPs) are the most abundant sequence variability in wheat genome. The nature of biallelic, cost-effective and high-throughput genotyping makes SNPs more suitable for genetic studies. The advent of wheat 90K iSelect SNP genotyping assay increased the number of gene-based markers which was applied for wheat genetic linkage map construction, genome-wide association studies and comparative genomics analysis (Cavanagh *et al.* 2013; Wang *et al.* 2014; Wu *et al.* 2015). In this study, *YrTZ2* was initially mapped into a 1.1 cM genetic interval between SSR markers *Xwmc230* and *Xgwm413*. To construct a high-density genetic linkage map, wheat 90K iSelect SNP genotyping assay was applied to saturate the genomic region of *YrTZ2*. Altogether, 250 polymorphic SNP markers clustering in 11 loci were located in the genomic region of *YrTZ2*. Finally, *YrTZ2* was delimited within a 0.8 cM genetic interval between locus *IWB19368* and marker *Xgwm413*, and co-segregated with locus *IWB28744* (consisting of 28 attaching SNP markers) that could be served as a starting point for chromosome landing and map-based cloning as well as marker-assisted selection (MAS) of the *YrTZ2* gene.

Comparative genomics analyses provided an effective way for wheat gene mapping. By applying comparative genomics analysis using genome sequences of *Brachypodium*, rice or sorghum, high-density genetic linkage maps of vernalization (*VRN*) genes (Yan *et al.* 2003, 2004, 2006), pairing homologous 1 (*Ph1*) (Griffiths *et al.* 2006), grain protein content-B1 (*Gpc*-*B1*) (Uauy *et al.* 2006), yellow rust resistance gene *Yr36* (Fu *et al.* 2009), wax production gene *W1* (Lu *et al.* 2015) and powdery mildew resistance gene *Pm6* (Qin *et al.* 2011), *Pm41 (*Wang *et al.* 2014), *Ml3D232* (Zhang *et al.* 2010), *MlIW170* (Liu *et al.* 2012; Liang *et al.* 2015), *MlIW172* (Ouyang *et al.* 2014) were constructed. The draft genome sequences of *T. aestivum* cv. Chinese Spring, *T. urartu* accession G1812 and *Ae. tauschii* accession AL8/78 enriched the available sequence resource and accelerated the wheat genomics research (Brenchley *et al.* 2012; Jia *et al.* 2013; Ling *et al.* 2013). The physical map of *Ae. tauschii*, anchored with 7,185 SNP marker extended sequence, provided an efficient tool for comparative genomics analyses among grass families, and marker development for fine mapping and map-based cloning of genes in wheat (Luo *et al.* 2013).

Comparative genomics analysis indicated highly collinearity between *YrTZ2* genomic region (*IWB19368*-*IWB31649*) of 1BS and a 24.5 Mb orthologous genomic region spanning 15 BAC-contigs of *Ae. tauschii* 1DS. The recently finished BAC-contig sequence of *Ae. tauschii* and Chinese Spring IWGSC whole genome assembly Ver. 1.0 would further contribute to fine mapping, map-based cloning and marker-assisted selection (MAS) of *YrTZ2*.

## ACKNOWLEDGEMENTS

This work was financially supported by the National Key Research and Development Program of China (2016YFD0101802).

---

**Table S1** Comparative genomics analysis among the *YrTZ2* locus, the genetic linkage map and physical map of *Aegilops tauschii*

## LITERATURE CITED

Akbari M., Wenzl P., Caig V., Carling J., Xia L., et al., 2006 Diversity arrays technology (DArT) for high-throughput profiling of the hexaploid wheat genome. Theor. Appl. Genet. %113: 1409–1420.

Akhunov E., Nicolet C., Dvorak J., 2009 Single nucleotide polymorphism genotyping in polyploid wheat with the Illumina GoldenGate assay. Theor. Appl. Genet. 119: 507–517.

Avni R., Nave M., Eilam T., Sela H., Alekperov C., et al., 2014 Ultra-dense genetic map of durum wheat×wild emmer wheat developed using the 90K iSelect SNP genotyping assay. Mol Breeding 34: 1549–1562.

Blanco A., Bellomo M. P., Cenci A., De Giovanni C., D’ovidio R., et al., 1998 A genetic linkage map of durum wheat. Theor. Appl. Genet. 97: 721–728.

Brenchley R., Spannagl M., Pfeifer M., Barker G. L., D’Amore R., et al., 2012 Analysis of the bread wheat genome using whole-genome shotgun sequencing. Nature 491: 705–710.

Cavanagh C. R., Chao S., Wang S., Huang B. E., Stephen S., et al., 2013 Genome-wide comparative diversity uncovers multiple targets of selection for improvement in hexaploid wheat landraces and cultivars. Proc. Natl. Acad. Sci. USA 110: 8057–8062.

Chagué V., Fahima T., Dahan A., Sun G. L., Korol A. B., et al., 1999 Isolation of microsatellite and RAPD markers flanking the *Yr15* gene of wheat using NILs bulked segregant analysis. Genome 42: 1050–1056.

Cheng P., Xu L. S., Wang M. N., See D. R., Chen X. M., 2014 Molecular mapping of genes *Yr64* and *Yr65* for stripe rust resistance in hexaploid derivatives of durum wheat accessions PI 331260 and PI 480016. Theor. Appl. Genet. 127: 2267–2277.

de Vallavieille-Pope C., Ali S., Leconte M., Enjalbert J., Delos M., et al., 2012 Virulence dynamics and regional structuring of *Puccinia striiformis* f. sp *tritici* in France between 1984 and 2009. Plant Dis. 96: 131–140.

Dvorak J., Terlizzi P., Zhang H. B., Resta P., 1993 The evolution of polyploid wheats: identification of the A genome donor species. Genome 36: 21–31.

Dvorak J., Zhang H. B., 1990 Variation in repeated nucleotide sequences sheds light on the phylogeny of the wheat B and G genomes. Proc. Natl. Acad. Sci. USA 87: 9640–9644.

Feldman M., 2001 The origin of cultivated wheat. In: Benjean AP, Angus J (eds) The wheat book: a history of wheat breeding. Lavoisier Publishing, Paris, pp 3-56

Fu D., Uauy C., Distelfeld A., Blechl A., Epstein L., et al., 2009 A kinase-START gene confers temperature-dependent resistance to wheat stripe rust. Science 323: 1357–1360.

Ganal M. W., Durstewitz G., Polley A., Berard A., Buckler E. S., et al., 2011 A large maize (*Zeamays* L.) SNP genotyping array: development and germplasm genotyping, and genetic mapping to compare with the B73 reference genome. PloS one 6: e28334.

Griffiths S., Sharp R., Foote T. N., Bertin I., Wanous M., et al., 2006 Molecular characterization of *Ph1* as a major chromosome pairing locus in polyploid wheat. Nature 439: 749–752.

Han D. J., Wang N., Jiang Z., Wang Q. L., Wang X. J., et al., 2012 Characterization and inheritance of resistance to stripe rust in the wheat line Guinong775. Hereditas 34: 1607–1613.

Jia J., Zhao S., Kong X., Li Y., Zhao G.,et al., 2013 *Aegilops tauschii* draft genome sequence reveals a gene repertoire for wheat adaptation. Nature 496: 91–95.

Krattinger S. G., Lagudah E. S., Spielmeyer W., Singh R. P., Huerta-Espino J., et al., 2009 A putative ABC transporter confers durable resistance to multiple fungal pathogens in wheat. Science 323: 1360–1363.

Li G. Q., Li Z. F., Yang W. Y., Zhang Y., He Z. H., et al., 2006 Molecular mapping of stripe rust resistance gene *YrCH42* in Chinese wheat cultivar Chuanmai 42 and its allelism with *Yr24* and *Yr26*. Theor. Appl. Genet. 112: 1434–1440.

Liang Y., Zhang D. Y., Ouyang S., Xie J. Z., Wu Q.H., et al., 2015 Dynamic evolution of resistance gene analogs in the orthologous genomic regions of powdery mildew resistance gene *MlIW170* in *Triticum dicoccoides* and *Aegilops tauschii*. Theor. Appl. Genet. 128: 1617–1629.

Lin F., Chen X. M., 2007 Genetics and molecular mapping of genes for race-specific all-stage resistance and non-race-specific high-temperature adult-plant resistance to stripe rust in spring wheat cultivar Alpowa. Theor. Appl. Genet. 114: 1277-1287.

Ling H. Q., Zhao S., Liu D., Wang J., Sun H., et al., 2013 Draft genome of the wheat A-genome progenitor *Triticum urartu*. Nature 496: 87–90.

Liu R. H., Meng J. L. 2003 MapDraw: a microsoft excel macro for drawing genetic linkage maps based on given genetic linkage data. Hereditas (Beijing) 25: 317–321.

Liu Z. J., Zhu J., Cui Y., Liang Y., Wu H. B., et al., 2012 Identification and comparative mapping of a powdery mildew resistance gene derived from wild emmer (*Triticum turgidum* var. *dicoccoides*) on chromosome 2BS. Theor. Appl. Genet. 124: 1041–1049.

Lu P., Qin J. X., Wang G. X., Wang L. L., Wang Z. Z., et al., 2015 Comparative fine mapping of the Wax 1 (*W1*) locus in hexaploid wheat. Theor. Appl. Genet. 128: 1595–1603.

Luo M. C., Deal K. R., Akhunov E. D., Akhunova A. R., Anderson O. D., et al., 2009 Genome comparisons reveal a dominant mechanism of chromosome number reduction in grasses and accelerated genome evolution in Triticeae. Proc. Natl. Acad. Sci. USA 106: 15780–15785.

Luo M. C., Gu Y. Q., You F. M., Deal K. R., Ma Y., et al., 2013 A 4-gigabase physical map unlocks the structure and evolution of the complex genome of *Aegilops tauschii*, the wheat D-genome progenitor. Proc. Natl. Acad. Sci. USA 110: 7940–7945.

Ma J. X., Zhou R. H., Dong Y. S., Wang L. F., Wang X. M., et al., 2001 Molecular mapping and detection of the yellow rust resistance gene *Yr26* in wheat transferred from *Triticum turgidum*L. using microsatellite markers. Euphytica 120: 219–226.

McIntosh R. A., Dubcovsky J., Rogers W. J., Morris C., Appels R., et al., 2014 Catalogue of gene symbols for wheat: 2013-2014 supplement. http://wwwshigennigacjp/wheat/komugi/genes/symbolClassListjsp

McIntosh R. A., Lagudah E. S., 2000 Cytogenetical studies in wheat. XVIII. Gene Yr24 for resistance to stripe rust. Plant Breed. 119: 81–83.

McIntosh R. A., Silk J., The T. T., 1996 Cytogenetic studies in wheat. XVII. Monosomic analysis and linkage relationships of gene *Yr15* for resistance to stripe rust. Euphytica 89: 395–399.

McIntosh R. A., Yamazaki Y., Dubcovsky J., Rogers J., Morris C., et al., 2013 Catalogue of gene symbols for wheat. 12th International Wheat Genetics SymposiumYokohama, Japan.

McIntosh R. A., Dubcovksy J., Rogers W. J., Morris C., Appels R., et al., 2016 Catalogue of gene symbols for wheat: 2015-2016 supplement.

Michelmore R. W., Paran I., Kesseli R. V., 1991 Identification of markers linked to disease-resistance genes by bulked segregant analysis: a rapid method to detect markers in specific genomic regions by using segregating populations. Proc. Natl. Acad. Sci. USA 88: 9828–9832.

Nachit M. M., Elouafi I., Pagnotta M. A., El Saleh A., Iacono E., et al., 2001 Molecular linkage map for an intraspecific recombinant inbred population of durum wheat (*Triticum turgidum* L. var. durum). Theor. Appl. Genet. 102: 177–186.

Ouyang S. H., Zhang D., Han J., Zhao X. J., Cui Y., et al., 2014 Fine physical and genetic mapping of powdery mildew resistance gene *MlIW172* originating from wild emmer (*Triticum dicoccoides*). PLoS One 9(6): e100160.

Peleg Z., Saranga Y., Suprunova T., Ronin Y., Roder M. S., et al., 2008 High-density genetic map of durum wheat × wild emmer wheat based on SSR and DArT markers. Theor. Appl. Genet. 117: 103–115.

Peng J. H., Fahima T., Röder M. S., Huang Q. Y., Dahan A., et al., 2000 High-density molecular map of chromosome region harboring stripe-rust resistance genes *YrH52* and *Yr15* derived from wild emmer wheat, *Triticum dicoccoides*. Genetica 109: 199–210.

Peng J. H., Fahima T., Röder M. S., Li Y. C., Dahan A., et al., 1999 Microsatellite tagging of stripe-rust resistance gene *YrH52* derived from wild emmer wheat, *Triticum dicoccoides*, and suggestive negative crossover interference on chromosome 1B. Theor. Appl. Genet. 98: 862–872.

Qin B., Cao A. Z., Wang H. Y., Chen T. T., You F. M., et al., 2011 Collinearity-based marker mining for the fine mapping of *Pm6*, a powdery mildew resistance gene in wheat. Theor. Appl. Genet. 123: 207–218.

Ramirez-Gonzalez R. H., Segovia V., Bird N., Fenwick P., Holdgate S., et al., 2015 RNA-Seq bulked segregant analysis enables the identification of high-resolution genetic markers for breeding in hexaploid wheat. Plant Biotechnol. J. 13: 613–624.

Riedelsheimer C., Lisec J., Czedik-Eysenberg A., Sulpice R., Flis A., et al., 2012 Genome-wide association mapping of leaf metabolic profiles for dissecting complex traits in maize. Proc. Natl. Acad. Sci. USA 109: 8872–8877.

Schneider A., Molnár I., Molnár-Láng M., 2008 Utilisation of *Aegilops* (goatgrass) species to widen the genetic diversity of cultivated wheat. Euphytica 163: 1–19.

Somers D. J., Isaac P., Edwards K., 2004 A high-density microsatellite consensus map for bread wheat (*Triticum aestivum* L.). Theor. Appl. Genet. 109: 1105–1114.

Song Q. J., Shi J. R., Singh S., Fickus E. W., Costa J. M., et al., 2005 Development and mapping of microsatellite (SSR) markers in wheat. Theor. Appl. Genet. 110: 550–560.

Sun G. L., Fahima T., Korol A. B., Turpeinen T., Grama A., et al., 1997 Identification of molecular markers linked to the *Yr15* stripe rust resistance gene of wheat originated in wild emmer wheat *Triticum dicoccoides*. Theor. Appl. Genet. 95: 622–628.

The International Wheat Genome Sequencing Consortium (IWGSC), 2014 A chromosome-based draft sequence of the hexaploid bread wheat (*Triticum aestivum*) genome. Science 345: 1251788.

Uauy C., Distelfeld A., Fahima T., Blechl A., Dubcovsky J., 2006 A NAC gene regulating senescence improves grain protein, zinc, and iron content in wheat. Science 314: 1298–1301.

Wan A. M., Chen X. M., He Z. H., 2007 Wheat stripe rust in China. Aust. J. Agr. Res. 58: 605–619.

Wan A. M., Zhao Z. H., Chen X. M., He Z. H., Jin S. L., et al., 2004 Wheat stripe rust epidemic and virulence of *Puccinia striiformis* f. sp *tritici* in China in 2002. Plant Dis. 88: 896–904.

Wang L. F., Ma J. X., Zhou R. H., Wang X. M., Jia J. Z., 2002 Molecular tagging of the yellow rust resistance gene *Yr10* in common wheat, P.I.178383 (*Triticum aestivum* L.). Euphytica 124: 71–73.

Wang S., Wong D., Forrest K., Allen A., Chao S., et al., 2014 Characterization of polyploid wheat genomic diversity using a high-density 90 000 single nucleotide polymorphism array. Plant Biotechnol. J. 12(6): 787–796.

Wang Z. Z., Cui Y., Chen Y. X., Zhang D. Y., Liang Y., et al., 2014 Comparative genetic mapping and genomic region collinearity analysis of the powdery mildew resistance gene *Pm41*. Theor. Appl. Genet. 127: 1741–1751.

Wu Q. H., Chen Y. X., Zhou S. H., Fu L., Chen J. J., et al., 2015 High-density genetic linkage map construction and QTL mapping of grain shape and size in the wheat population Yanda 1817 × Beinong6. PLoS One 10(2): e0118144.

Xie W. L., Nevo E., 2008 Wild emmer: genetic resources, gene mapping and potential for wheat improvement. Euphytica 164: 603–614.

Yan L., Fu D., Li C., Blechl A., Tranquilli G., et al., 2006 The wheat and barley vernalization gene *VRN3* is an orthologue of *FT*. Proc. Natl. Acad. Sci. USA 103: 19581–19586.

Yan L., Loukoianov A., Blechl A., Tranquilli G., Ramakrishna W., et al., 2004 The wheat *VRN2* gene is a flowering repressor down-regulated by vernalization. Science 303: 1640–1644.

Yan L., Loukoianov A., Tranquilli G., Helguera M., Fahima T., et al., 2003 Positional cloning of the wheat vernalization gene *VRN1*. Proc. Natl. Acad. Sci. USA 100: 6263–6268.

Zhang H. T., Guan H. Y., Li J. T., Zhu J., Xie C. J., et al., 2010 Genetic and comparative genomics mapping reveals that a powdery mildew resistance gene *Ml3D232* originating from wild emmer co-segregates with an NBS-LRR analog in common wheat (*Triticum aestivum* L.). Theor. Appl. Genet. 121: 1613-1621.

Zhang J. Y., Xu S. C., Zhang S. S., Zhao W. S., Zhang J. X., 2001 Monosomic analysis of resistance to stripe rust for source wheat line Jinghe8811. Acta. Agron. Sin. 27: 273–277.

Zhao K., Tung C. W., Eizenga G. C., Wright M. H., Ali M. L., et al., 2011 Genome-wide association mapping reveals a rich genetic architecture of complex traits in *Oryza sativa*. Nat. Commun. 2: 467.

